# *NLMR* and *landscapetools*: An integrated environment for simulating and modifying neutral landscape models in R

**DOI:** 10.1101/307306

**Authors:** Marco Sciaini, Matthias Fritsch, Cédric Scherer, Craig Eric Simpkins

**Affiliations:** Department of Ecosystem Modelling, University of Göttingen, Büsgenweg 4, 37077, Göttingen, Germany; Department of Ecological Dynamics, Leibniz Institute for Zoo and Wildlife Research, Alfred-Kowalke-Str. 17, 10315, Berlin, Germany

**Keywords:** artificial pattern, landscape generator, neutral landscape model, R, spatial visualisation, virtual landscape

## Abstract

1. Neutral landscape models (NLMs) simulate landscape patterns based on theoretical distributions and can be used to systematically study the effect of landscape structure on ecological processes. NLMs are commonly used in landscape ecology to enhance the findings of field studies as well as in simulation studies to provide an underlying landscape. However, their creation so far has been limited to software that is platform dependent, does not allow a reproducible workflow or is not embedded in R, the prevailing programming language used by ecologists.
2. Here, we present two complementary R packages *NLMR* and *landscapetools*, that allow users to generate, manipulate and analyse NLMs in a single environment. They grant the simulation of the widest collection of NLMs found in any single piece of software thus far while allowing for easy manipulation in a self-contained and reproducible workflow. The combination of both packages should stimulate a wider usage of NLMs in landscape ecology. *NLMR* is a comprehensive collection of algorithms with which to simulate NLMs. *landscapetools* provides a utility toolbox which facilitates an easy workflow with simulated neutral landscapes and other raster data.
3. We show two example applications that illustrate potential use cases for *NLMR* and *landscapetools:* First, an agent-based simulation study in which the effect of spatial structure on disease persistence was studied. Here, spatial heterogeneity resulted in more variable disease outcomes compared to the common well-mixed host assumption. The second example shows how increases in spatial scaling can introduce biases in calculated landscape metrics.
4. Simplifying the workflow around handling NLMs should encourage an uptake in the usage of NLMs. *NLMR* and *landscapetools* are both generic frameworks that can be used in a variety of applications and are a further step to having a unified simulation environment in R for answering spatial research questions.

## Introduction

Neutral landscape models (NLMs) are algorithms which generate landscape patterns in the absence of specific biotic and abiotic processes (*Gardner et al.*, 1987; Li *et al.*, 2004; Caswell, 1976). These models were originally developed as null models used to test landscape scale hypotheses (Gardner & Urban, 2007; With & King, 1997). NLMs are now used in a wide variety of ways to examine and test observations and metrics of ecological patterns and processes at landscape scales (With & King, 1997; *Turner et al.*, 2001). With & King (1997) and Turner & Gardner (2015) defined three uses for NLMs: (i) coupling ecological models with NLMs to predict the change of ecological processes, (ii) analyse the extent of structural deviation between real and neutral landscapes, and (iii) development and testing of new landscape metrics. The large number of applications and their increased uptake has led to the development of a number of different software designed to generate NLMs, using a range of algorithms. Researchers can use standalone software such as RULE (Gardner, 1999), its successor QRULE (Gardner & Urban, 2007) and SIMMAP (Saura & Martínez-Millán, 2000), as well as LG (van Strien *et al.*, 2016). However, in our experience the most useful pieces of software are those that offer a collection of different NLM algorithms based in an open source programming language and allowing for a reproducible and consistent workflow. Packages developed for common programming languages, such as NLMpy (*Etherington et al.*, 2015) for the *PYTHON* coding environment, fulfill those criteria. However, currently there is no NLM software package available for the programming language most widely used by ecologists, the *R* programming language (R Core Team, 2017). Examining 100 recent articles published in *Methods in Ecology and Evolution* we found a clear trend in the community towards *R* with 70 publications utilizing R (see supplementary materials, Fig. 1). The lack of a flexible framework in R may limit the overall use of NLMs. We have, therefore, developed the two complementary R packages, *NLMR* and *landscapetools*, to allow for the easy simulation and manipulation of NLMs and other rasters.

**Figure 1:**
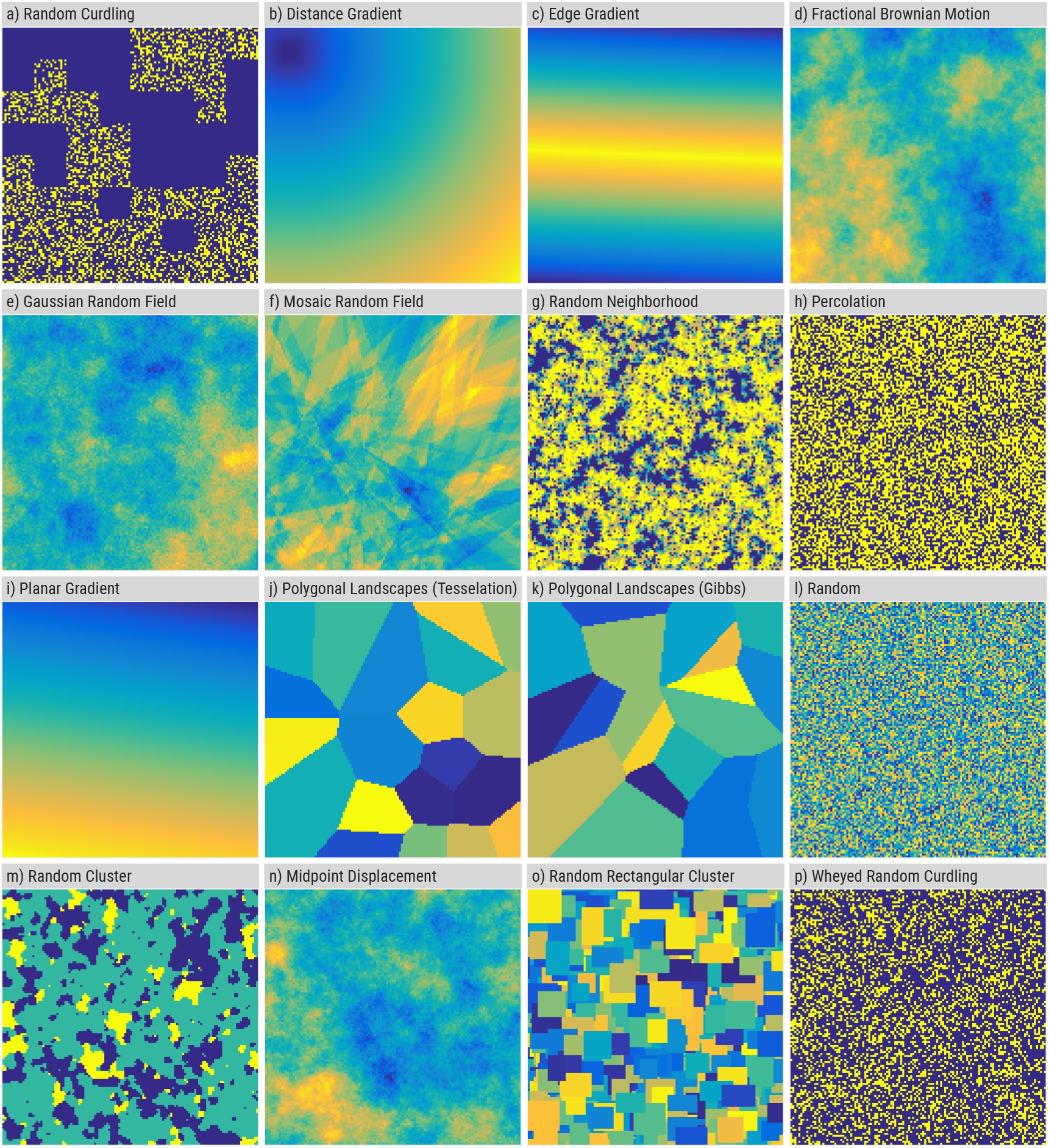
Collection of all neutral landscape models (NLM) implemented in the *NLMR* package (Table 1). See section Data accessibility for R code to create each of the NLMs.

Both packages will allow a growing group of scientific R users to make use of NLMs in their research while permitting for a streamlined workflow contained in the R environment. Furthermore, the integration of the major geographical information system libraries (such as GDAL and GEOS) in R embed our packages in a framework that allows every analysis and simulation to be done in R without relying on proprietary software.

## Functionality

*NLMR* and *landscapetools* both build on the established geospatial environments in R, e.g. the *raster* (Hijmans, 2017) and *sf* packages (Pebesma, 2018). The nomenclature of all our functions was designed to facilitate a reproducible workflow while being distinct from the naming conventions of other geospatial packages, thus avoiding namespace conflicts with other functions and packages.

## NLMR

The *NLMR* package is a generic numeric framework to generate NLMs using the widest collection of algorithms found in any single piece of software, while also enabling for the combination and integration of different NLM algorithms. All NLM functions in *NLMR* (Table 1) simulate two-dimensional raster objects. By default, no spatial reference system is applied but can be incorporated, allowing NLMs to be projected to the spatial extent of any study area. Algorithms differ from each other in terms of spatial autocorrelation, ranging from no autocorrelation (i.e. random NLM) to a constant gradient (i.e. planar gradients) (Fig. 1).

**Table 1:**
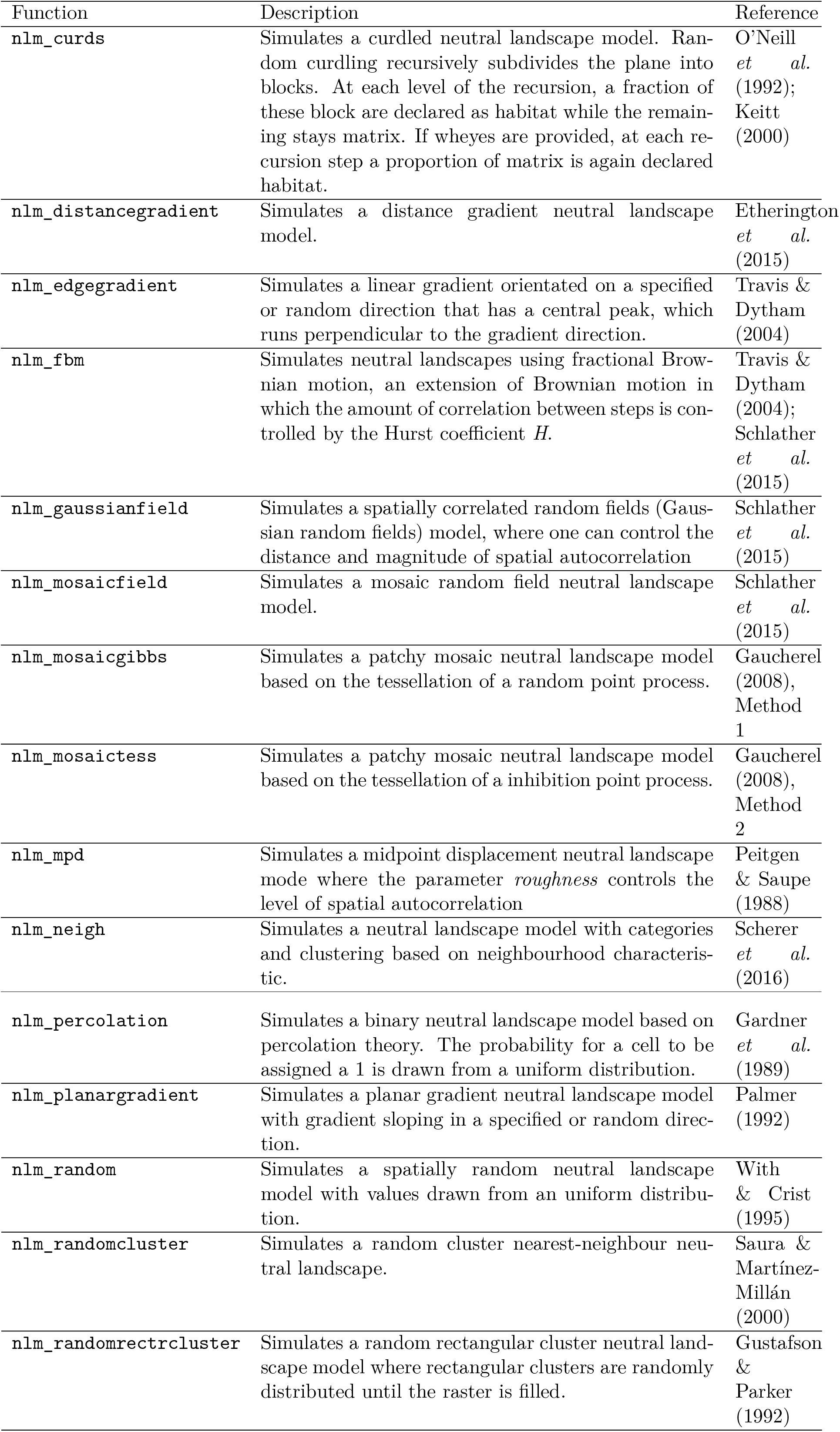
List of implemented neutral landscape models in NLMR with a short description and the associated literature for more information.

The basic syntax used to simulate a NLM is: `nlm_modeltype(ncol, nrow, resolution, …)`

The raster objects returned from NLMR can readily be transformed and visualised by *landscapetools* or incorporated in spatially-explicit analyses and simulation models.

### landscapetools

The package *landscapetools* contains a set of utility function to complete tasks involved in most landscape analysis (Fig. 2). This includes visualisation, (reclassification and merging methods thus providing a workflow to apply NLM algorithms in a broad landscape ecology context. All the functions in *landscapetools* require a raster object as an input.

**Figure 2:**
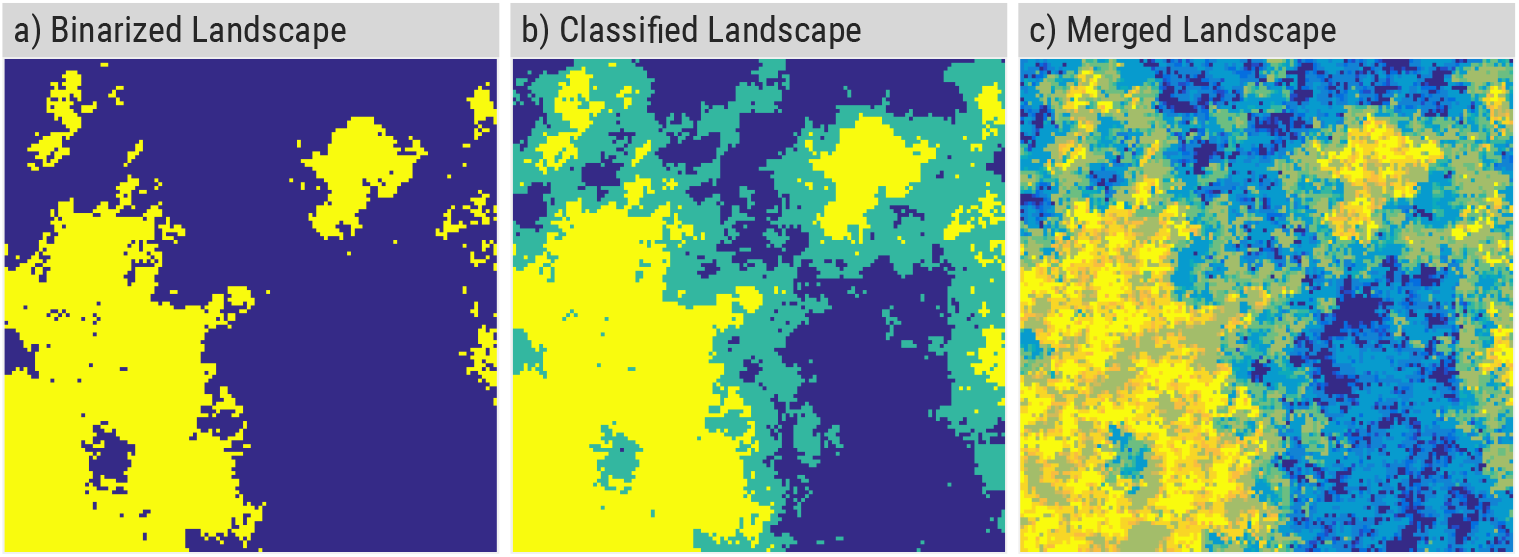
Visualisation of the main functionality of *landscapetools* with an underlying fractional brownian motion neutral landscape model (`nlm_fbm()`). a) shows the `util_binarize()` function, that classifies landscapes into habitat and matrix. b) is the same landscape classified (`util_cassify()`) into three landscape categories with equal proportions. c) shows the the classified landscape from b) merged (`util_merge()`) with a random neighbourhood neutral landscape model (Fig. 1g)). See section Data accessibility for R code about how to use *landscapetools.*

The functions `util_classify and util_binarize` (re-)classify raster data into proportions based upon given weightings. `util_classify` has auxiliary functions that simplify the use of factorial encoding of landscape data in a raster object. `util_binarize` returns landscapes with a binary representation of the raster, e.g. matrix/habitat distributions. As simulation experiments often rely on multiple differing proportions of matrix and habitat, it is possible to define multiple breaks and compute a collection (RasterStack) of binarized landscapes at once. The function `util_merge` allows a weighted combination of multiple landscape rasters thus allowing for even more sophisticated landscape patterns. The combination of NLMs is an established way of deriving ecotones where the focus is on merging planar gradients with less autocorrelated landscapes (Travis & Dytham, 2004; *Etherington et al.*, 2015). `util_rescale` is internally used in all algorithms implemented in NLMR, but is also a public function in *landscapetools* to linearly rescale raster cell data into a range between zero and one.

*landscapetools* offers functions and themes for the clear visual communication of spatial data as *ggplot2* objects (Wickham, 2009). The visualisation focuses on achieving outputs that can be used in the context of publications, consequently applying a clear theme with the colorblind friendly color scales from the viridis package (Garnier, 2018) and typographic elements that support a reproducible workflow. Further auxiliary functions in *landscapetools* help to coerce raster data in tibbles (Müller & Wickham, 2018), the new standard for rectangular data in R, and vice versa.

## Example applications

This section describes two example applications of *NLMR* and *landscapetools* (code available in the section Data accessibility): i) investigate disease persistence in heterogeneous landscapes and ii) estimate bias resulting from increased scaling of landscape patterns.

### Case study 1: Disease persistence in heterogeneous landscapes

Simulation models often assume underlying landscapes to be homogeneous or simply suitable habitat distributed in an unsuitable matrix. Given the comprehensive collection of NLMs with differing amounts of autocorrelation provided by *NLMR*, we are able to test the effect of homogeneous versus heterogeneous underlying landscapes on ecological dynamics. We demonstrate the effect of spatial heterogeneity on the dynamics of a directly transmitted disease, Classical Swine Fever, infecting a wild boar population. We use a modified version of an existing agent- and grid-based simulation model (*Kramer-Schadt et al.*, 2009; *Lange et al.*, 2012). Transmission between individuals is calculated based on the health status of individuals in the same and adjacent cells.

Heterogeneous landscapes (50 × 25 cells) were simulated using the function `nlm_mosaictess()` with two levels of fragmentation (low fragmentation with *germs* = 25 and high fragmentation with *germs* = 250). Total carrying capacities were equal for all simulated landscapes. Patchy, fragmented mosaic neutral landscapes were selected to represent typically fragmented landscape structures in which the boars are located. The flexibility of the NLM algorithm allowed for different levels of fragmentation to be easily simulated. *landscapetools* allowed for the classification of the generated landscapes into ecologically reasonable breeding capacities per cell and analysis of our results, all within the R environment. Our results show that disease outcomes are less variable in homogeneous setups compared to heterogeneous landscapes which might underestimate predictions of disease outcomes (Fig. 3).

**Figure 3:**
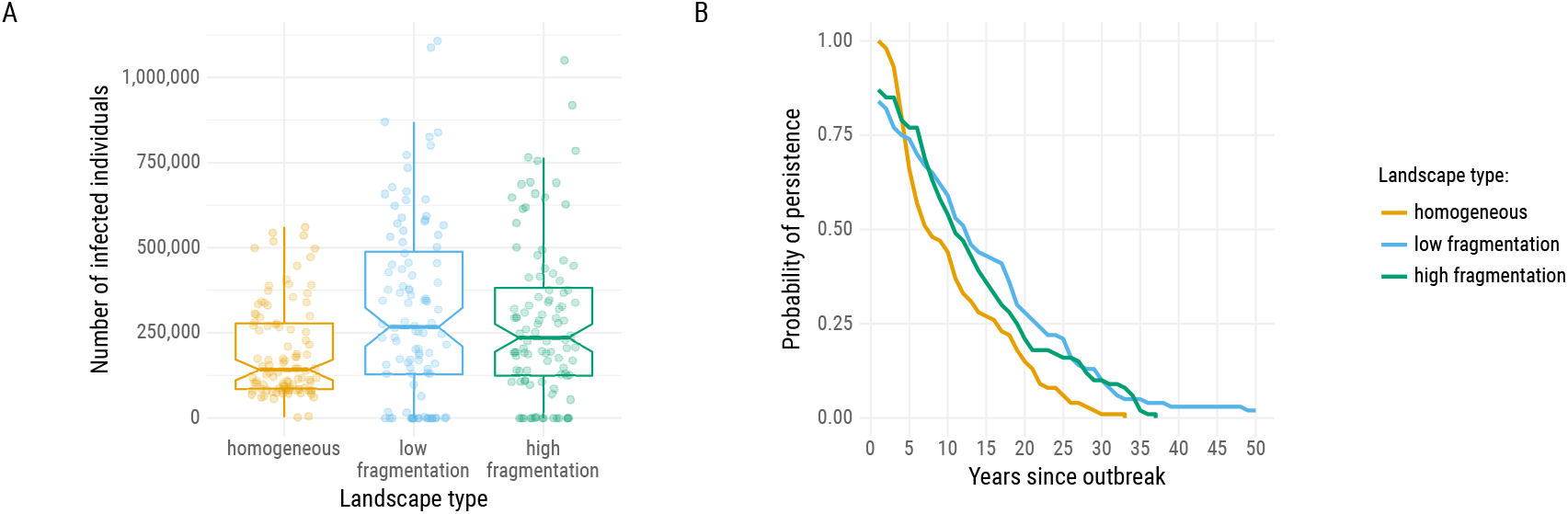
Disease outcomes using different landscape types in an epidemiological model. Landscape type “low fragmentation” describes a pattern of some large patches (`nlm_polylands(gem = 25)`) while landscapes with “high fragmentation” consist of multiple small patches of varying host capacities (`nlm_polylands(gem = 250)`). (A) Overall number of infected individuals per landscape type shown as boxplot and raw data. (B) Probability of the disease being persistent over time, estimated as yearly ratio of simulations with ongoing infections per overall simulations (100 per landscape type).

### Case study 2: Bias estimation when scaling up landscape patterns

Altering the resolution of a map can result in a systematic bias of metrics describing any contained spatial patterns. We utilize *NLMRs’* functionality to generate differing neutral landscapes as a solid base for investigating this phenomenon. Initially, we started with landscapes similar to the ones used by Bocedi et al. (2012).

To highlight several possible scaling outcomes we implemented two different aggregation methods.

First, we generated multiple landscapes (1024 × 1024 cells) with the desired amount of habitat patches (p = 0.1, 0.3, 0.5, 0.7) and fragmentation grade (H = 0.1, 0.5, 0.9) by using *NLMRs* midpoint displacement algorithm (`nlm_mpd()`). Second, we used a simple averaging and majority rule as aggregation methods to scale up the landscape (Fig. 4). Third, we calculated the desired landscape metric *p* (proportion of suitable cells) for each scaling step and visualise possible biases (Fig. Supp. 2).

**Figure 4:**
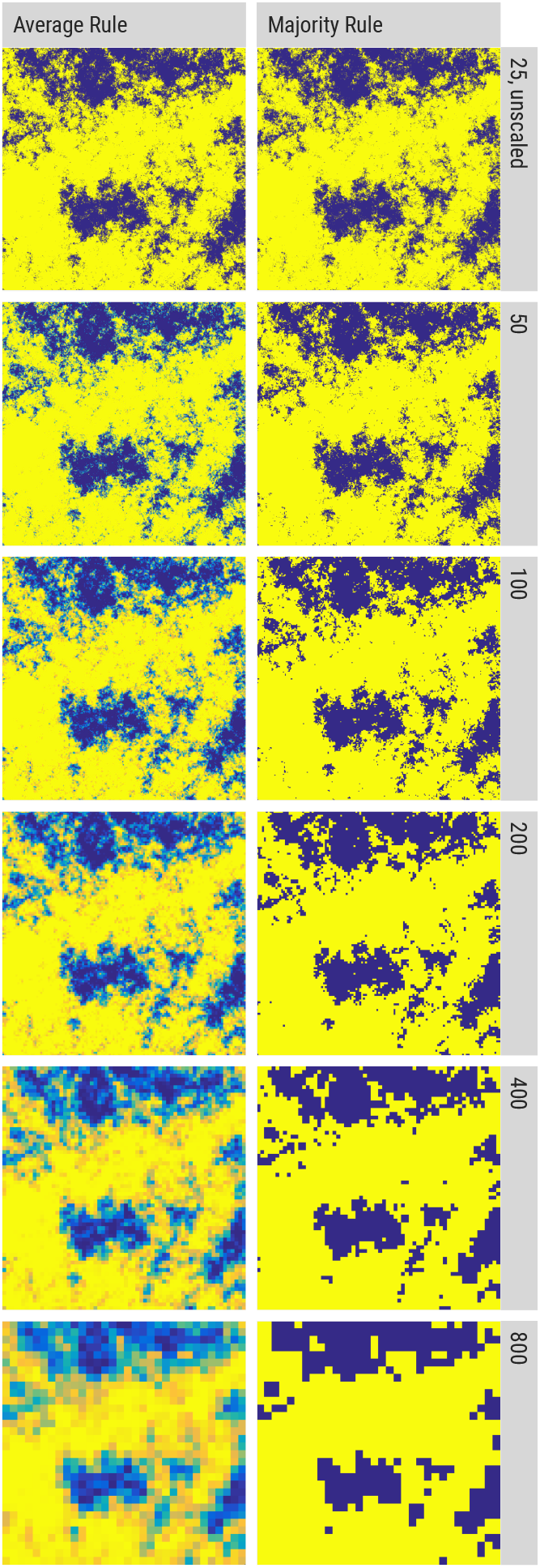
Visualisation of the aggregation methods used in case study 2: Aggregation by average rule (left column) and by majority rule (right column). Cell size range from 25 m to 800 m. The landscape example was generated with *p* = 0.7 and *H* = 0.9. The underlying neutral landscape model is the midpoint displacement algorithm (simulated with `nlm_mpd()` and `util_binarize()`).

This simple example demonstrates the systematic over-and underestimation of the targeted metric that could happen when using scaling methods, depending on the initial landscape.

## Conclusion and Outlook

*NLMR* is the first software to allow the simulation of NLMs within the selfcontained, reproducible framework of R. It can be applied in all landscape analyses in which one wants to test the influence of NLMs on ecological dynamics. Although existing tools are capable of simulating some of the NLMs contained in *NLMR* (Gardner, 1999; Gardner & Urban, 2007; Saura & Martínez-Millán, 2000; *van Strien et al.*, 2016; *Etherington et al.*, 2015), none of them combines as many different types and none of them are as well integrated in a native geospatial workflow. Hence, the majority of the limitations that previous NLM software exhibited, such as developing own methods for spatial operations like masking and extracting, are overcome. The functions in *landscapetools* provide a variety of different established methods to handle landscape data. We believe that our functions are supplementary to packages like *rasterVis* (Perpiñán & Hijmans, 2018), which enhance the visualising of landscape patterns allowing for the clear communication of landscape research.

*NLMR* and *landscapetools* are both designed to simplify the workflow of scientific landscape analyses. Thus, it is extensively documented and the implementation of each simulation model is based on published literature. Every function provides examples in its respective help file. A vignette in *NLMR* combines the information about the functions and the examples into an introduction of a basic workflow for using NLMs. Future extensions of *NLMR* and *landscapetools* aim to include all established algorithms that have been used to create NLMs and accompanying utility functions.

We believe that being capable of simulating NLMs natively in R is highly beneficial for the field of landscape ecology. *NLMR* and *landscapetools* have been officially released on CRAN (current versions 0.3.0) and peer-reviewed by rOpen-Sci (https://github.com/ropensci/onboarding/issues/188).

## Acknowledgements

MS was supported by the German Research Foundation, DFG, through grant number WI 1816/18-1 as part of the DFG Research Unit FOR2432/1 and as part of the Scaling Problems in Statistics Research Training Group DFG-GRK 1644/3. MF was supported by the German Research Foundation, DFG, as part of the Scaling Problems in Statistics Research Training Group DFG-GRK 1644/2. CS was supported by the German Research Foundation, DFG, as part of the BioMove Research Training Group DFG-GRK 2118/1. We thank Kerstin Wiegand for her help in improving the manuscript. We thank rOpenSci for their extensive code review.

## Authorship statement

M.S. and C.E.S. conceived of the original package concept. M.S. led code development, and all other coauthors were co-developers. M.S. drafted the manuscript, with case studies analysed and drafted by C.S. and M.F. All coauthors contributed critically to the drafts and gave final approval for publication.

## Data accessibility

The *NLMR* package and documentation are hosted at https://github.com/ropensci/NLMR. The *landscapetools* package and documentation are hosted at https://github.com/ropensci/landscapetools. Reproducible scripts for the figures and case studies are available from https://github.com/marcosci/Sciaini_et_al_2018.

## Supplementary materials

**Supplementary 1:**
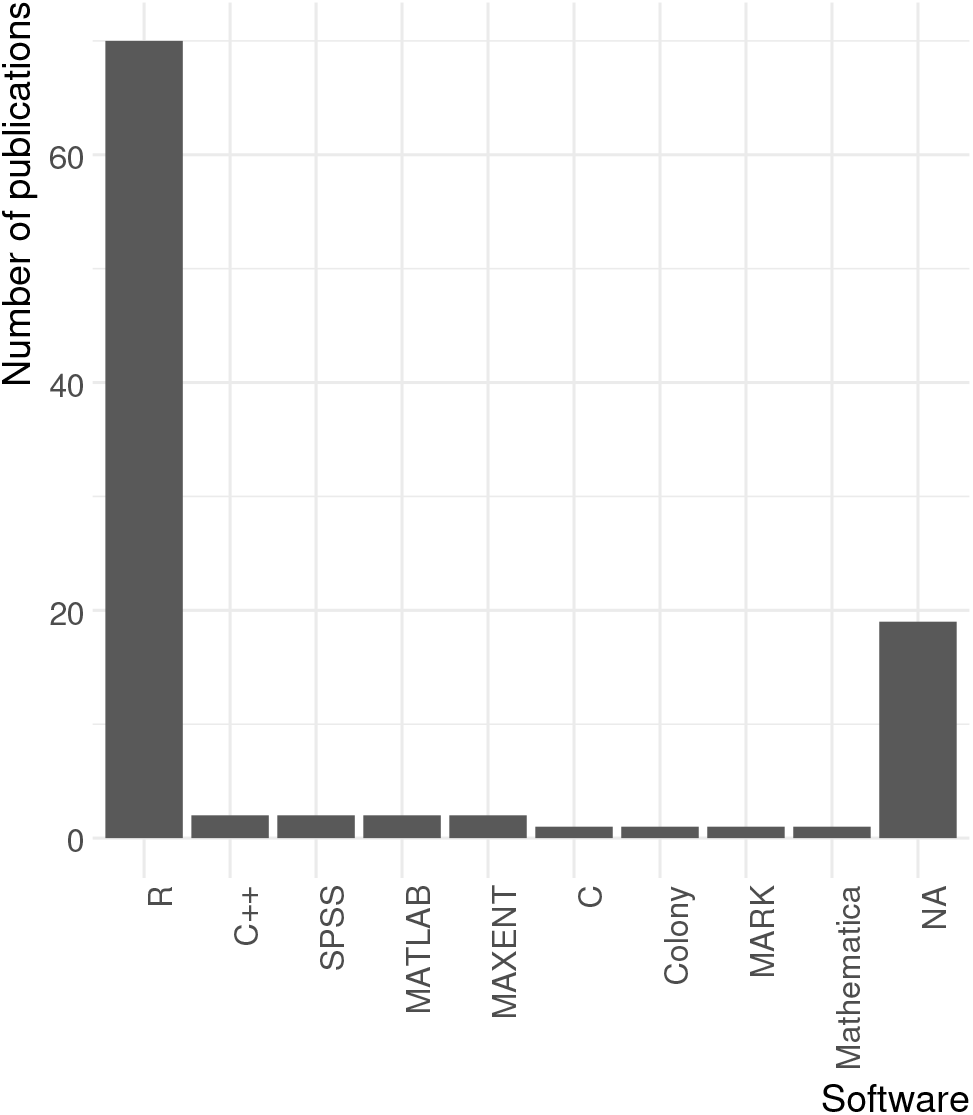
Software usage from 100 recent publications from Methods in Ecology and Evolution in 2017. If there was no information on what type of software was used we classified it as NA.

**Supplementary 2:**
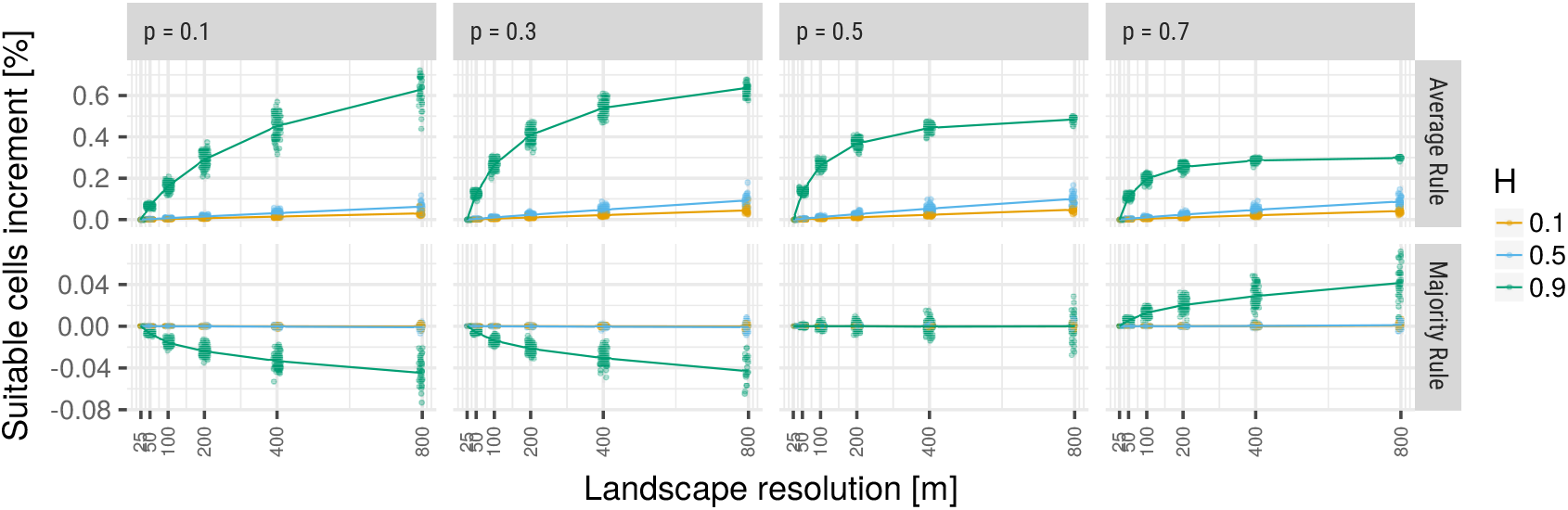
Mean increment of suitable cells in the landscapes. Each dot is the absolute difference from the scaled up landscape to the original one. The line visualises the mean at each scaling step. Negative values and means indicate underestimation, positive ones indicate overestimation of suitable cells. Increasing values of *p* indicate an increasing amount of habitat patches and the higher *H* the higher the fragmentation grade of the landscape structure.

